# Retrovirus insertion site analysis of LGL leukemia patient genomes

**DOI:** 10.1101/535997

**Authors:** Weiling Li, Lei Yang, Robert S. Harris, Lin Lin, Thomas L. Olson, Cait E. Hamele, David J. Feith, Thomas P. Loughran, Mary Poss

## Abstract

**Background:** Large granular lymphocyte (LGL) leukemia is an uncommon cancer characterized by a sustained clonal proliferation of LGL cells. Antibodies reactive to retroviruses have been documented in the serum of patients with LGL leukemia. Culture or molecular approaches have to date not been successful in identifying a retrovirus.

**Methods:** Because a retrovirus must integrate into the genome of an infected cell, we focused our efforts on detecting a novel retrovirus integration site in the clonally expanded LGL cells. We present a new computational tool that uses long-insert mate pair sequence data to search the genome of LGL leukemia cells for retrovirus integration sites. We also utilize recently published methods to interrogate the status of polymorphic human endogenous retrovirus type K (HERV-K) provirus in patient genomes.

**Results:** While our analysis did not reveal any new retrovirus insertions in LGL genomes from LGL leukemia patients, we did identify four HERV-K provirus integration sites that are polymorphic in the human population and absent from the human reference genome, hg19. To determine if the prevalence of these or other polymorphic proviral HERV-Ks differed between LGL leukemia patients and the general population, we applied a recently developed approach that reports all sites in the human genome occupied by a proviral HERV-K. Using the 1000 genomes project (KGP) data as a reference database for HERV-K proviral prevalence at each polymorphic site, we show that there are significant differences in the number of polymorphic HERV-Ks in the genomes of LGL leukemia patients of European origin compared to individuals with European ancestry in the KGP data.

**Conclusions:** Our study confirms that the integration of a new infectious or endogenous retrovirus does not cause the clonal expansion of LGL cells in LGL leukemia, although we do not rule out that these cells could be responding to retroviral antigens produced in other cell types. However, it is of interest that the burden of polymorphic proviral HERV-K is elevated in LGL leukemia patient genomes. Our research emphasizes the merits of comprehensive genomic assessment of HERV-K in cancer samples and suggests that further analyses to determine contributions of HERV-K to LGL leukemia are warranted.

## Background

Large granular lymphocyte (LGL) leukemia is a rare, chronic, proliferative disorder of cytotoxic T cells (approximately 85% of cases) and NK cells [1]. Diagnosis of this leukemia is based on a sustained elevation of a clonally expanded T or NK cell population. LGL leukemia is reported most frequently in patients from North America and Europe and up to half of patients also have an autoimmune disorder, most frequently rheumatoid arthritis [2]. A small subset of patients show clonal proliferation of CD4+ cells, which has been associated with Cytomegalovirus infection [3]. An aggressive form of LGL leukemia involving natural killer cells, (NK leukemia) is most common in East Asians and has been linked with Epstein-Barr virus infection [4]. Aspirates demonstrate close approximation of LGL and antigen presenting cells [5], emphasizing that prolonged presentation of an unknown antigen could be a common underlying feature of the various forms of LGL leukemia. At present, it is unknown if an infectious agent is responsible for chronic antigen stimulation of LGL in some or all patients, although non-malignant proliferation of LGL does occur in chronic viral infections [6].

Serum antibodies from LGL leukemia patients recognize an antigen with homology to a protein encoded by human T-lymphotropic virus (HTLV) [7, 8], providing a potential link of this disease with retroviruses. The oncogenic potential of retroviruses is well established in mammals and birds, which can develop cancers of hematopoietic cells [9] following retroviral infection. Although there are numerous mechanisms for retrovirus-induced oncogenesis, dysregulation of key cell cycle control genes during retroviral integration and transduction of cellular oncogenes are particularly well documented [9–11]. While many cancers in animals have a retrovirus etiology, HTLVs are the only retrovirus group definitively linked to cancer in humans. In this case, a virus-encoded accessory protein is necessary for cell transformation [12, 13]. Despite a high prevalence of HTLV-1 antigen responders among LGL leukemia patients, intensive molecular and culture-based approaches have failed to detect HTLV, or any known retrovirus, in LGL leukemia patients. However, such methods could fail to identify a defective retrovirus, which is relevant because many animal oncogenic retroviruses are replication-defective and can have unusual genome sequence [14, 15]. Because LGL leukemia involves a clonal expansion of LGL cells, we reasoned that if a retrovirus initiated the malignancy, we should be able to detect the integrated virus in LGL genomes even if it couldn’t be recovered in culture. The goal of this study was to interrogate LGL genomes of LGL leukemia patients for the presence of a novel retrovirus to determine if the clonal expansion of LGL cells is preceded by the integration of a novel or known retrovirus.

## Methods

### Patient samples

T-LGL leukemia patients met the clinical criteria of T-LGL leukemia with increased numbers of CD3+, CD8+/CD57+ T lymphocytes or CD3^-^, CD16+/CD56+ NK cells in the peripheral blood[16]. LGL leukemia patient blood samples were obtained and informed consents signed for sample collection according to the Declaration of Helsinki using a protocol approved by the Institutional Review Board of the University of Virginia. Blood was subjected to Ficoll-Hypaque (Sigma Aldrich) gradient centrifugation for peripheral blood mononuclear cell (PBMC) isolation.

### Whole genome sequencing (WGS)

Long insert mate pair libraries of 11 LGL patients (S1-S11) were prepared and sequenced at the Duke Center for Genomic and Computational Biology using the Illumina Nextera MP kit. Sequencing was performed on a HiSeq and read length was 125bp. Paired-end sequencing of 48 LGL patients was conducted by Illumina. Patients S9, S10, and S11 were paired-end sequenced at Penn State by Dr. Stephan Schuster.

### PCR assessment of four non-reference polymorphic HERV-K

A 25 μl PCR reaction was performed using human adult normal peripheral blood leukocyte genomic DNA purchased from Biochain (Cat. #D8234148-1, Lot #B511221) and the following primer pairs:

Chr1_5HK For (5′CATAGCAAATCCCAGTGTAGACATC3′) and

Chr1_5HK Rev (5′CTGGGAGCATTTCTGGACATC3′);

Chr1_PREINT For (5′CACCGCACCTGGCAAGTTTACA3′) and

Chr1_PREINT Rev (5′ATTTGGGGTCCTCATGAAGCAGAA3′);

Chr19_3HK5 For (5′TACCCCAAGACCAAAAATAATAAG3′) and

Chr19_3HK6 Rev (5′CTGATAGTGGCAAGATGGATGTA3′);

C19PRE3 For (5′GAACAGGAGCATGCTCATAGTGTGT3′) and

C19PRE4 Rev (5′GGTCTCGAACTCCTAACCTCCTG3′);

Chr12PREINT For (5 ′TAC T GGGAAT AAGAT GAT GAT GGT3′) and

Chr12PREINT Rev (5′TTTGTTAAGTGCTCGGAAGGT3′);

Chr10HK1 For (5′ACGCGTGGGTATATCGGTTTATTTCT3′) and

Chr10HK2 Rev (5′GGCTGGTTCTTTATTATTTATGGCTGGT3′);

Chr12HK5_1 For (5′GCAGTTCCACCTTCCCGACAGC3′) and

Chr12HK5_2 Rev (5′ACACAGGACCAAAAGAACGAGT AATC3′).

Each 25 μl reaction contained 12.5 μl 2x MyTaq HS Red Mix, 2 μl 5μM forward primer, 2 μl 5μM reverse primer, 3.5 μl nuclease-free water and 20 ng of the pre-dispensed genomic DNA. The reactions were cycled using a Bio-Rad C1000 Touch 96-deep well cycler. The cycled reaction and HyperLadder 1 kb DNA ladder were electrophoresed on a 1% agarose gel with 1x TAE for 60 minutes at 80 V. The gel was visualized using 10,000x Sybr Safe DNA Gel Stain and images were captured using the Bio-Rad ChemiDoc and Image Lab software.

### Insertion Call Pipeline

A detailed description of the insertion call pipeline is given in Additional File S1_Supplementary Methods Methods. Briefly, sites containing a retroviral insertion result in aberrant mapping of WGS mate pairs to the reference genome. We developed a series of signal tracks to detect insertions that include shorter than expected insert length of mate pairs, distant or inter-chromosomal mapping of mate pairs, orphan mate pairs where only one read of the pair can be mapped, and partially mapped reads (Figure 1). Parameters of the caller were tuned using a simulation to detect 5-12 kbp insertions of at least 55% prevalence in a sample. The tracks were integrated to identify candidate insertion events. The sequence content of each candidate insertion was investigated by gathering the read pairs that map near the insertion, assembling them and querying the resultant contigs against the NCBI nt database using BLAST. BLAST results were searched for infectious or endogenous retrovirus hits based on taxonomy identification (Figure S3). Endogenous retrovirus hits that had host flanking regions were confirmed to be from the candidate insertion site by mapping to the reference genome. Polymorphic HERV-K insertions were confirmed with an approach (Additional File S1_Supplementary Methods Methods and [17]) independent of the insertion call pipeline.

**Figure 1.**
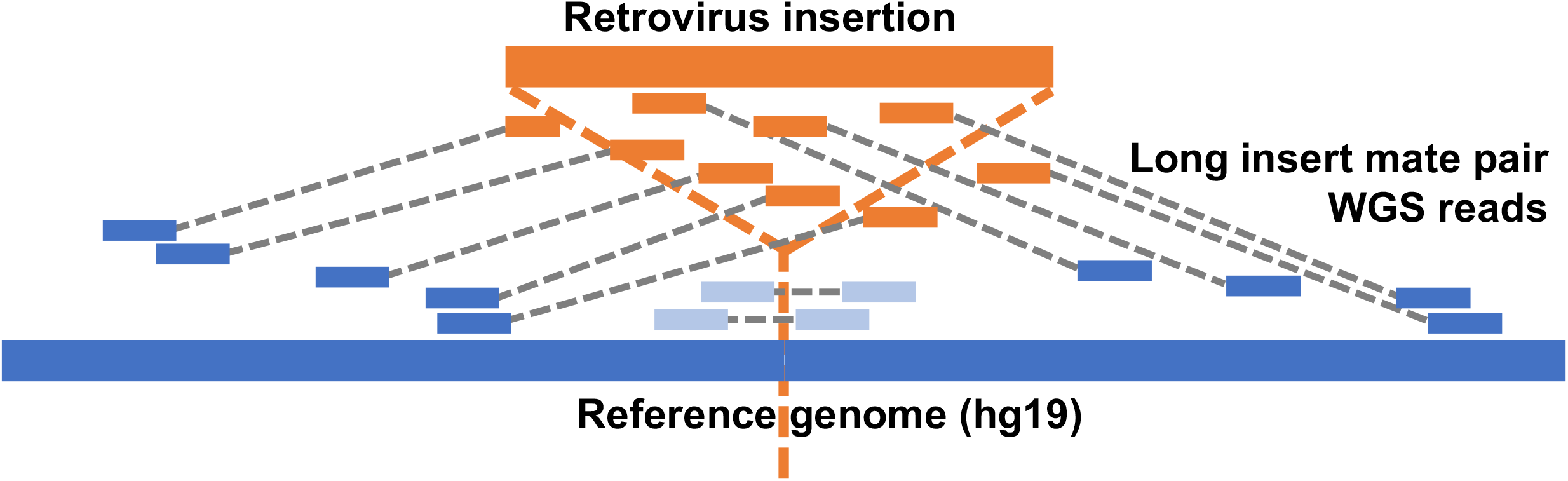
Utilizing long insert mate pair reads to localize retrovirus integrations. Reference human genome is shown as a blue line with the location of an inserted retrovirus, in orange, indicated by a dotted vertical orange line. Long insert mate pair reads are linked by gray dotted lines, with the read derived from the new retrovirus, which will not map, shown in orange, it’s mate that maps to the human reference genome shown in blue; and mate pairs that both map to the reference with a short insert length shown in light blue. A retrovirus insertion site is suggested by a combination of several features of mate pair mapping including short insert intervals and discordant or broken mate pairs. The unmapped reads (orange in the figure) from discordant mate pairs at each called insertion site are assembled and used to determine the sequence of a candidate retrovirus.

## Results

### Detecting retroviral integrations in LGL leukemia patient genomes

There is compelling evidence for retroviral involvement in LGL leukemia [18, 19]. Because a retrovirus must integrate into the host genome as part of its life cycle, we focused our methodology to detect retroviral insertions in the genomes of LGL cells from LGL leukemia patients because they are a clonal cell population. Identifying a unique retrovirus integration site in a patient’s genome is complicated because reads derived from the retrovirus will not map to a reference genome. Thus, we developed a two-step pipeline (Additional File S1_Supplementary Methods) that first detects insertion events in the patient genome and then determines if the insertion is of retroviral origin. Both steps of our method exploit specific mapping properties of long insert (8 kbp) mate pair data to identify retrovirus insertions (Figure 1).

Briefly, a read within approximately 5-12 kbp of a newly inserted retrovirus will map to the reference genome (anchoring mate, dark blue in Figure 1) while its mate will be derived from the retrovirus and will not map (unmapped mate, orange in Figure 1) or could map to a location such as an endogenous retrovirus element elsewhere in the genome if there is sufficient homology (discordant mate); either case results in a broken pair of read mates with one mapping to the reference genome and one that does not map within the appropriate distance. Depending on the length of the retrovirus, which typically is 6-10 kbp, some mate pairs may span the entire inserted virus and hence both mate pairs will originate from the host (light blue in Figure 1). However, because the retrovirus is not present in the reference genome, these mate pairs will map at a distance shorter than the expected insert distribution of 5-12 kbp. The insert length and depth of mapped reads are key signals in our retrovirus insertion pipeline.

We used a simple simulation to explore how features of long insert mate pair mapping could be integrated to call an insertion compatible with a retroviral integration, to set parameters, and to determine how the pipeline performed for retroviruses of different lengths and at different prevalence in the sample. (Additional File S1_Supplementary Methods, Figure S1 and Table S1). The results of the simulation demonstrated that our ability to detect a retrovirus insertion depended both on the length of the retrovirus and the prevalence in the sample. If more than 80% of the cells carry a retroviral integration of any length, they are detectable in the simulation. Retroviruses longer than 8 kbp cannot reliably be detected if their prevalence in the sample is less than 60%. This is primarily because we lose the signal from the short insert length track with longer retrovirus insertions because longer viruses will cause the insert length to exceed that represented by our mate pair library and only one of the mates will map. We account for longer retroviruses in an alternative approach (discussed below, see Additional File S1_Supplementary Methods). The proportion of LGL in the peripheral blood of LGL leukemia patients is typically 65-80%, which is within the range that our simulation indicates we can reliably detect retroviruses across the size spectrum. Thus, we applied the pipeline to our WGS data, using the parameters established by the simulation.

We first applied this pipeline to the long insert mate pair WGS data of LGL leukemia patient S10, who had a PBMC count consisting of 80% clonal LGL cells. These data are displayed as tracks for a genome browser and each candidate insertion site can be accessed from a catalogue of all insertion calls. The first step of the insertion call pipeline is intentionally permissive to false positives because we implement a second step that utilizes the discordant mate pair data to detect retroviral sequences at each called insertion. Discordant mates identified at each called insertion site (represented in Track 4, Figure S1 and S2) are assembled and queried against the NCBI database using BLAST+ [20, 21] (Additional File S1_Supplementary Methods). We anticipated that the majority of insertions would be known genome structural variants in the human population that are not in the hg19 reference genome but are represented in the NCBI nt database. If this is the case, multiple contigs generated from assembling the discordant mate pairs at the called insertion will have a best match (lowest e-value) to the same database entry, and have a taxonomy ID of ‘human’. A taxonomy ID and name search of the BLAST output using ‘retrovirus’ was applied to all contigs that were not identified as human. There were no detectable retrovirus matches in S10 sequence data using these criteria.

We further queried candidate insertions for key word ‘endogenous retrovirus’ (ERV), because there are polymorphic ERVs in the human population that are absent from hg19, many of which are represented in BAC clones; these would also appear as an insertion in our pipeline. Candidate human endogenous retrovirus (HERV) insertions were confirmed by two criteria: the contigs derived from unmapped mates mapped to a HERV in the NCBI nt database and the host regions flanking the HERV in the NCBI nt entry could be aligned to the reference human genome (hg19) in the interval defined by the anchoring mates near the called insertion. We detected four polymorphic HERV-K elements absent from hg19 in patient S10; these include two sites (chr1:73594980-73595948; chr10:27182399-27183380) containing a solo LTR in hg19, one site that had been previously reported as polymorphic (chr12:5727215-55728183) and one site that was recently reported to be polymorphic [22] (chr19:22414379-22414380), all of which we confirmed empirically. The pipeline was then used to analyze 10 additional LGL leukemia patient samples (S1-S9 and S11). None of the LGL patients had an unknown retrovirus sequence detectable in the DNA of PBMC but all patients had the polymorphic HERV-K provirus at chr1:73594980-73595948 and chr10:27182399-27183380, while nine had the HERV-K at chr12:5727215-55728183 and five at chr19:22414379-22414380.

Our data indicate that there is no clonal integration of a novel retrovirus, either exogenous or endogenous, in the LGL leukemia cells. However, this does not rule out that a retrovirus contributes to this disease. Our samples come from PBMC and spleen, both of which contain normal cells that are frequently targets for retrovirus infection. The insertion call tool was designed to detect a retrovirus integration of 9 kbp or less in the clonally expanded LGL cells; a longer retrovirus or a retrovirus that integrated into non-leukemic cells would be below the level of sensitivity of our method. We took two additional approaches to search for a low frequency integration event in the sample. All unmapped reads that passed a quality filter were assigned taxonomy identification provided from SNAP [23] mapping (Additional File S1_Supplementary Methods Methods). All reads with a ‘virus’ taxonomy classification were further scrutinized by BLAST search to determine if the best match was to a retrovirus. None were identified. We also mapped all long insert mate pair WGS reads to a full length HERV-K (GenBank accession number: JN675087) and assigned their mates to a position in hg19 to search for candidate de novo insertions of a HERV-K. Again, none were detected. Thus, we can confirm that there is not a clonal integration of an unknown exogenous or endogenous retrovirus in the LGL leukemia cells themselves and that we were not able to detect evidence of a novel retrovirus integration site in the genomes of non-leukemic cells that were represented in our WGS data.

This investigation was motivated by data showing that LGL patients have sero-recognition of HTLV proteins, although they are not infected with this virus [8, 18, 24]. Our detailed investigation of LGL genomes from 11 leukemia patients (S1-S11) failed to detect a novel retrovirus integration site in circulating LGL that could elicit this antibody response, but we did identify several HERV-Ks that are polymorphic in humans and absent from the human reference genome. Endogenous retroviruses have been implicated in several cancers including leukemia [25, 26] and immune response to HERV proteins has been reported in both cancer and autoimmune diseases [27–29]. Of the reported polymorphic HERV-K, 16 are close to full length [17, 22]. However, the distribution of all polymorphic HERV-K in cancer patients compared to the population at large is at present unknown. We thus investigated whether the prevalence of polymorphic HERV-K - individually or co-occurring - or the total number of polymorphic HERV-K in a person’s genome differed between LGL leukemia patients and normal populations represented in the KGP data [17, 30].

Our detection method to identify retroviral insertions used in this paper depended on long insert mate pair sequencing, which is not typically available for the large genomic databases needed to determine the prevalence of polymorphic HERV-K in global populations. We recently reported on a method that utilizes unique k-mers present in each published HERV-K provirus to estimate the proviral prevalence of polymorphic HERV-K proviruses in any individual from paired-end sequence data [17]. The output of the pipeline is the ratio *(n/T)* of k-mers from a query set (n) to the total number of unique k-mers (T) for each HERV-K proviral insertion. We applied this approach to a set of 51 LGL leukemia patients, 11 of which (S1-S11) also had long-insert mate pair sequence data available.

As previously noted [17], the distribution of polymorphic HERV-K proviruses varies considerably among KGP populations. Most LGL leukemia patients are of European descent and have clonal proliferation of T cells although our sample includes eight patients with LGL leukemia involving NK cells and three individuals of non-European ancestry. Forty of the patients with T-LGL leukemia are of European origins, therefore we present the data both for all 51 LGL patients versus KGP and for the 40 T-LGL-EUR patients compared to European KGP data (EUR). Our analyses include 90 fixed and polymorphic HERV-K proviruses, omitting three on the Y chromosome, those recently reported to be expanding in centromeres [31, 32], and chr1:73594980-73595948 [17]. The provirus frequency for both the entire LGL patient population (51 individuals) and the 40 T-LGL-EUR falls within the range of values for the five KGP populations for all HERV-K proviruses except those at chr1:75842771-75849143, chr12:58721242-58730698 and chr3:148281477-148285396, where prevalence in LGL patients is higher than any of the KGP populations (Table 1, see Table S2 for the full data set); and chr19:21841536-21841542 and chr19:22414379-22414380, where LGL patients have a lower prevalence than any of the five KGP populations. chr12: 58721242-58730698 is noteworthy because 98% of T-LGL-EUR patients carry this HERV-K compared to 87% of EUR, which is the highest of the five KGP populations.

**Table 1.** Prevalence (proportion) of LGL patients and individuals from the five super-populations represented in the KGP data carrying a polymorphic HERV-K. AFR: African, AMR: Admixed American, EAS: East Asian, EUR: European, SAS: South Asian, LGL: All LGL patients in this study (51 total), ELGL: T-LGL patients with European ancestry (T-LGL-EUR, 40 total).

| HERV-K (hg19 coordinate) | LGL | ELGL | KGP | AFR | AMR | EAS | EUR | SAS |
| --- | --- | --- | --- | --- | --- | --- | --- | --- |
| **chr1:75842771-75849143** | 70.6 | 70.0 | 42.9 | 26.8 | 56.5 | 6.0 | 68.9 | 66.8 |
| **chr3:148281477-148285396** | 49.0 | 52.5 | 41.9 | 38.9 | 42.6 | 45.0 | 46.5 | 37.4 |
| **chr12: 58721242-58730698** | 94.1 | 97.5 | 70.7 | 58.9 | 78.4 | 60.0 | 87.3 | 75.5 |
| **chr19: 21841536-21841542** | 5.9 | 7.5 | 27.0 | 39.2 | 11.9 | 32.2 | 10.7 | 32.4 |
| **chr19:22414379-22414380** | 51.0 | 47.5 | 67.8 | 89.2 | 60.8 | 56.9 | 55.8 | 67.2 |

We verified the estimates of HERV-K presence in LGL patients from our pipeline in two ways. The four polymorphic HERV-K proviruses that are absent in hg19 were identified in our insertion call pipeline and we confirmed the presence of a provirus using discordant mate pair sequences for 11 patients (S1-S11). We also used the mate pair data to confirm the status of the remaining 16 polymorphic HERV-K that are represented in hg19; the results for HERV-K status based on our data mining tool and mate pairs are 100% concordant. In addition, we amplified both the preintegration site and a portion of HERV-K including the host flanking sequence for the four polymorphic HERV-K that were identified in the insertion call pipeline in 48 of the LGL patients (LGL) and 48 individuals with no diagnosed diseases (European, African-American and Hispanic origin; hereafter referred to as “normal”). The results from the PCR assay for chr19:22414379-22414380 (normal 56%, LGL patient 50%) and chr12:55727215-55728183 (normal 82%, LGL patients 71%) agree with our computational analysis (Table S2). The chr10:27182399-27183380 HERV-K was amplified in all LGL patients and normal individuals, which is consistent with the high prevalence (99%) found in our data mining method. chr1:73594980-73595948 was also present in all patients and normals by PCR but we have no data on this virus from our KGP analysis because the build of the reference genome (GRCh37) used to map KGP reads included hs37d5, a concatenated decoy sequence which contains this virus, while our approach used coordinates of hg19. Hence the reads needed to identify chr1:73594980-73595948 were not extracted in the data mining step [17].

We performed a linear discriminant analysis (LDA) to determine if the signal in the HERV-K prevalence data was sufficient to distinguish the T-LGL-EUR patient population from EUR. For this analysis we used only the 28 individuals from the KGP data sets with high coverage (~30x sequencing depth) data, after confirming that none were outliers in their population clusters of all KGP data (~5x sequencing depth) [17]. Based on the data reduced to the states ‘absence, solo LTR, provirus’ of each HERV-K insertion, T-LGL-EUR patients separate from EAS and AFR but were admixed with EUR, AMR and SAS (Figure 2A). We previously showed that using the *n/T* ratio provided better resolution of the KGP populations [17], presumably because it captures allelic differences in both fixed and polymorphic HERV-K. Based on *n/T,* T-LGL-EUR patients are well separated from all KGP populations (Figure 2B). These data indicate that both the polymorphic HERV-K and specific allelic forms of each HERV-K provirus define the T-LGL patient population.

**Figure 2.**
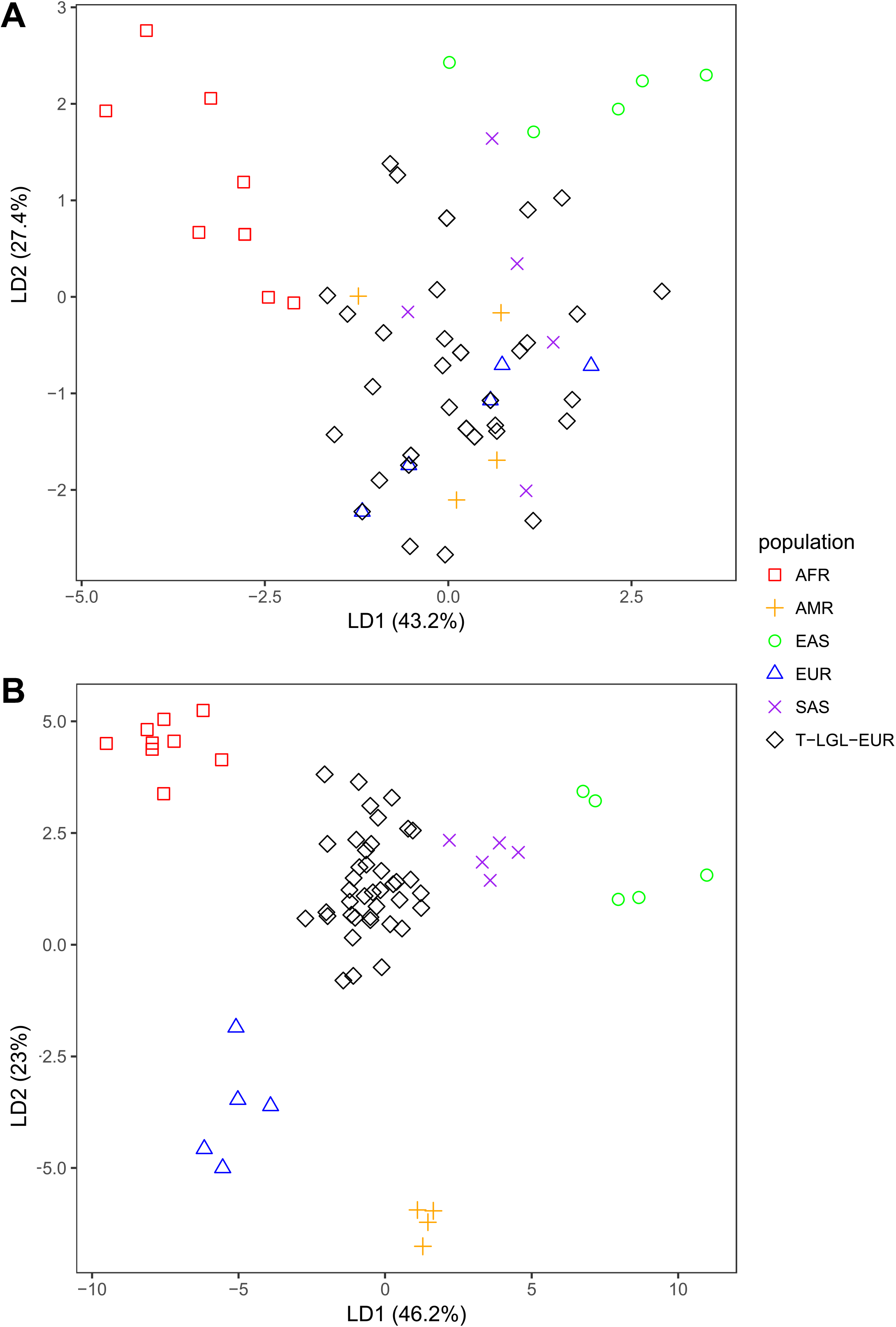
Linear discriminant analysis based on HERV-K status of T-LGL-EUR patients and KGP super populations. Linear discriminant analysis (LDA) was conducted on data generated by a comprehensive analysis of polymorphic HERV-Ks in an individual genome [17] A. Data is based on three HERV-K states of ‘absence, solo LTR, provirus’ or B. the *n/T* ratio of each known HERV-K provirus for T-LGL leukemia patients of European ancestry and the 28 individuals from KGP super populations with high coverage data. The ratio indicates the proportion of k-mers derived from a person’s WGS dataset (n) that are 100% match to a set of unique k-mers (T) characterizing each HERV-K provirus. The improved resolution of T-LGL-EUR patients from other individuals using n/T likely reflects that alleles of HERV-K contribute to population differentiation. The symbols and colors for each KGP populations and T-LGL-EUR leukemia patients are indicated in the key on the right.

An *n/T* ratio of 1 indicates that the reference allele (typically from hg19) is present. We suggested in [17] that *n/T* less than 1 indicates that an individual has an allele of the HERV-K at a locus that is not represented in the database. Using the unmapped mates from those reads flanking a HERV-K locus, we reconstructed the sequence of a polymorphic HERV-K proviruses at chr3:112743479-112752282, which is presented here because there is considerable variation in *n/T* in both LGL patients (Figure S4A) and KGP [17]. For individuals with n/T=1, all reads map perfectly to the unique k-mers representing the reference alleles. However, LGL patients with *n/T<1* have five substitutions in this HERV-K, one common to the 11 patients (S1-S11) with long insert mate pair data and four sites that were variably present among these individuals (Figure S4B), which accounts for the differences in *n/T.* These data demonstrate that *n/T* can reflect allelic differences at a HERV-K locus and indicate why the *n/T* ratio contains more information than the presence, absence data to distinguish populations.

Genomic structural variations are often noted in cancer cells of diverse origins [33, 34]. Because HERV-Ks are polymorphic in the genome some individuals have a higher burden of these repetitive elements than others. We considered that an increased number of polymorphic HERV-K proviruses could contribute to the sustained clonal proliferation that characterizes LGL leukemia by increasing chromosomal re-arrangements [35–37]. In the KGP datasets, no individual had fewer than 7 or more than 18 of the 20 polymorphic HERV-K proviruses evaluated and ~50% of all individuals from each of the KGP populations have 12 or 13 polymorphic provirus insertions except for EAS, where 52% of the sampled individuals have 10 or 11 polymorphic integration sites [17]. T-LGL-EUR patients carry between 9 and 16 HERV-K proviruses (Figure 3) at proportions that are significantly different than those found in EUR individuals (Kolmogorov-Smirnov test, p=0.0087; Table S2). Notably, 35% of T-LGL-EUR individuals carry 14 proviruses while the carriage rate for this number of HERV-K proviruses among the five KGP populations is 2-22%. Hence, the LGL leukemia patients sampled in this analysis carry a higher burden of polymorphic HERV-K than is seen in the general population.

**Figure 3.**
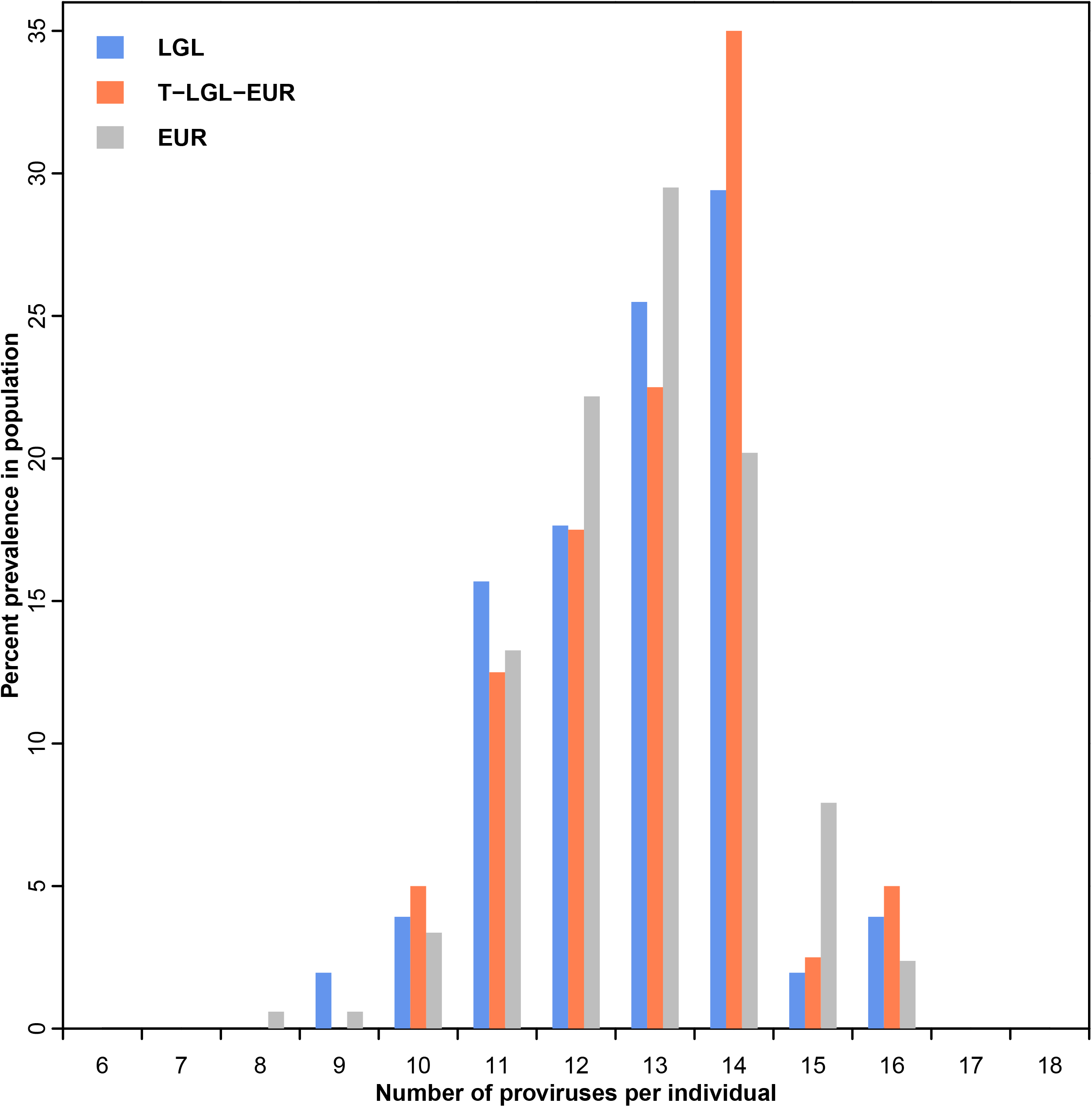
Histogram of the number of polymorphic HERV-K proviruses identified in LGL leukemia patients compared to individuals of European origin from KGP. Data are shown for 51 LGL patients (blue) and for the subset of 40 patients with T-LGL leukemia of European ancestry (T-LGL-EUR, orange). Data for the 505 EUR individuals (gray) from the KGP data is from Li *et al.* [17].

Because LGL leukemia patients, and particularly T-LGL leukemia patients, have more proviruses in their genomes, we reasoned that co-occurrence of the polymorphic HERV-Ks could also vary from EUR or other global populations. This is the case for several HERV-K combinations that include chr12: 58721242-58730698, which is present in 98% of T-LGL leukemia patients (Figure 4A). Interestingly, there are combinations of HERV-K, with or without chr12: 58721242-58730698, that are higher in LGL patients than in EUR but similar to AFR and EAS populations, which have the lowest prevalence of chr12: 58721242-58730698 of the KGP populations (Figure 4B). These data suggest that the prevalence of co-occurring HERV-K should be considered when investigating a role of HERV-K in the pathogenesis of a specific disease.

**Figure 4.**
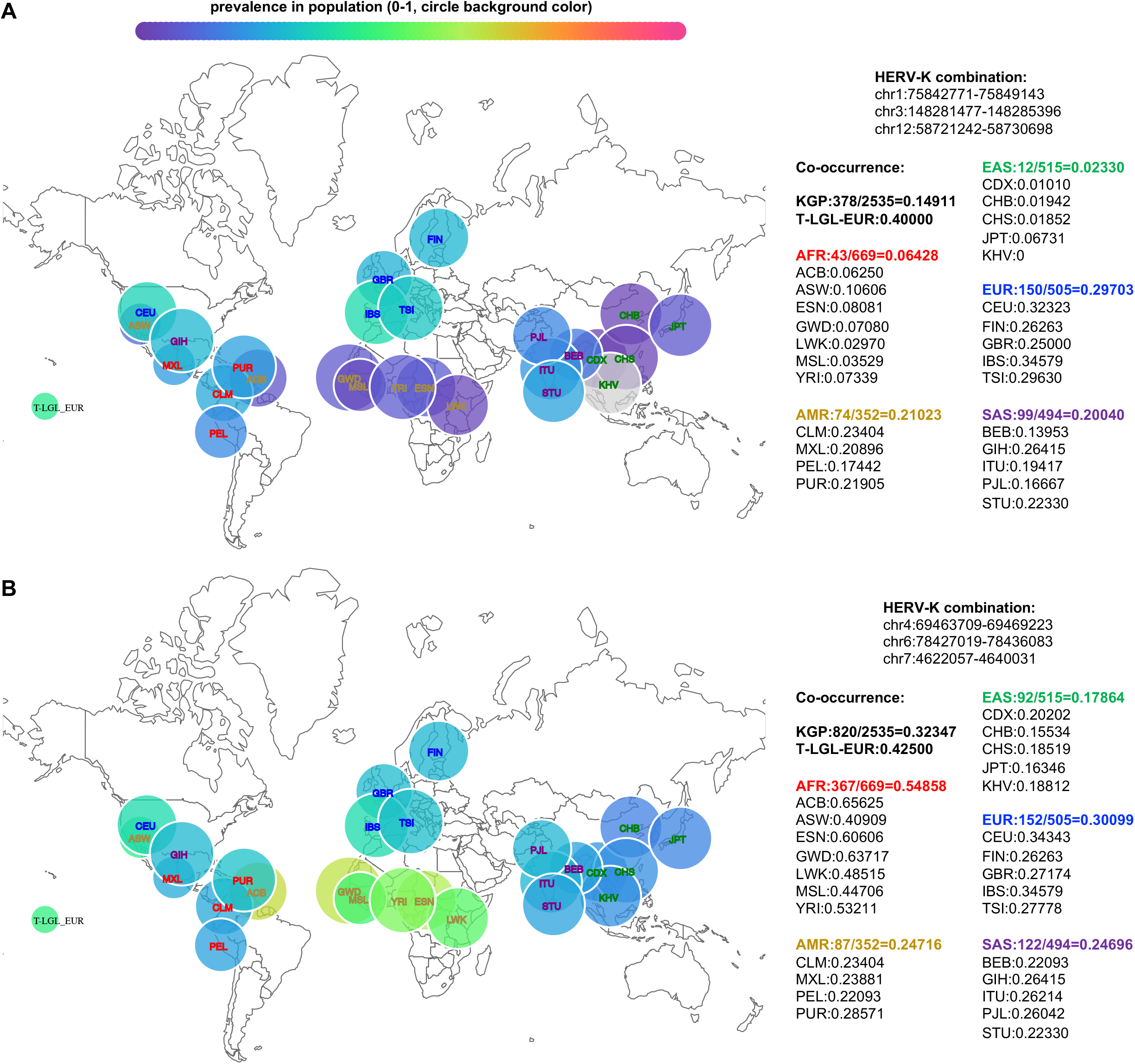
The prevalence of combinations of polymorphic HERV-K provirus in KGP populations and T-LGL-EUR leukemia patients. The combinations of polymorphic HERV-K provirus evaluated are indicated at the top right of each panel. **A**. The prevalence of three polymorphic HERV-K proviruses that include chr12: 58721242-58730698 in KGP individuals and T-LGL-EUR patients. **B**. The prevalence of three polymorphic HERV-K, excluding chr12: 58721242-58730698, in KGP individuals and T-LGL-EUR leukemia patients. Coordinates are referenced to hg19. Bubble size is proportional to the number of individuals in the population and color gradient represents prevalence from 0 to 100%. Absolute values are given in the text on the right for each population. KGP population abbreviations are given in Table 1 and additional information can be found at (http://www.internationalgenome.org/category/population/).

## Discussion

Our goal was to determine if the serological reactivity to retroviral antigens reported in LGL leukemia patients could be explained by the integration of a novel retrovirus in LGL genomes. Insertional mutagenesis is a common mechanism of cell transformation, and is in part a consequence of where the retrovirus integrates in the genome; proximity to a host gene can result in both altered regulation of the gene and expression of the retrovirus [38, 39]. If retrovirus integration causes the clonal expansion of LGL, the insertion site should be present in the genome of leukemic cells from LGL leukemia patients. We developed a tool to detect retrovirus insertion sites based on long insert mate pair data and conclude that there is not a new retroviral integration site in the clonally expanded cells. However, our tool revealed several polymorphic HERV-Ks in the LGL leukemia patient genomes that are not present in the human reference genome. Hence, we applied additional methods that we recently reported [17] to investigate the genome wide distribution of HERV-K in LGL leukemia patients compared with unaffected individuals represented in the KGP data.

The most notable difference between LGL leukemia patients and individuals represented in KGP data is in the burden of polymorphic HERV-K proviruses that they carry. This difference is more pronounced when restricting the comparisons to only the 40 individuals with T-LGL leukemia who are of European ancestry (T-LGL-EUR). This likely reflects the fact that EAS populations represented in KGP have a significantly lower overall burden of polymorphic HERV-K and there are three LGL leukemia patients of East Asian descent in our test cohort. LGL patients also have an elevated prevalence of chr12:58721242-58730698 and several combinations of polymorphic HERV-K are found more frequently in LGL patients than in EUR individuals, although not all involve the chr12:58721242-58730698 HERV-K.

Our previous analysis of the KGP data also suggested that there were alleles of HERV-K proviruses not found in the NCBI databases; these are represented by an *n/T* ratio of less than 1 [17]. We reconstructed the provirus sequences at HERV-K loci using patient long insert mate pair WGS data to confirm that our analysis tool does report allelic differences in HERV-K that are not found in any of the reference HERV-K that localize to that site. We call these unknown alleles because we require 100% match of query k-mers from patient WGS to the set of k-mers, T, which represent k-mers unique to all alleles present in public databases of a HERV-K at a specific locus. If a k-mer derived from a patient contains a sequence polymorphism at any position of the unique reference k-mer set T, it will be excluded in the k-mer count, effectively decreasing *n/T* to less than 1. It is notable that there is substantial variation in *n/T* for both fixed and polymorphic HERV-K and these differences, not presence or absence of a HERV-K, distinguish the KGP super-populations [17]. Although population-specific alleles have been reported [40], our data highlight more extensive sequence variation among HERV-Ks than has previously been recognized and suggest that both sequence and occupancy of a site should be considered when assessing the potential role of HERV-K in disease. This is an important consideration because using a consensus sequence or specific reference sequence might not reflect the HERV-K in a patient population.

Our study was motivated by serological evidence of a retrovirus in LGL patients that was not a known human retrovirus. The data we present do not rule out the contribution of an infectious retrovirus to LGL leukemia pathogenesis. The insertion call pipeline developed herein would only identify a clonal integration of a new retrovirus in the LGL leukemia cells, which comprise greater than 60% of the peripheral mononuclear cell population, or a new somatic or germline insertion in the case of a polymorphic HERV-K. Our conclusion that the cells sampled from patients do not harbor an integrated retrovirus is additionally supported by the negative results of the SNAP analysis, which interrogated all reads for evidence of a retrovirus. Thus, we consider several possibilities to account for the serological response of patients and the absence of data indicating that the responding cells are infected. Cells infected with a retrovirus in any tissue could express viral antigen without producing infectious particles that could stimulate an antibody response; serological reactivity could also be targeted to aberrant expression of a HERV-K [25]. Given the well-established potential of infectious retroviruses to activate and recombine with ERVs [30, 41–46], an additional and intriguing consideration is that a chimeric, replication-incompetent retrovirus exists in LGL leukemia patients. These possibilities may be investigated by further immunological analyses to understand the nature of the antigens that can either induce an anti-retroviral response or sustain proliferation of LGL or both.

## Conclusions

Our results indicate that LGL leukemia patients have a genomic landscape of polymorphic HERV-K that is different than populations at large. Thus, a thorough analysis of sequence, structural and epigenetic variation in proximity to individual and co-occurring polymorphic HERV-K may reveal if HERV-K contribute to the pathogenesis of this leukemia. Because a considerable number of LGL leukemia patients also have an autoimmune disease, further comprehensive investigation of the roles of endogenous or exogenous retroviruses in LGL leukemia and autoimmune disease is indicated.

## Abbreviations

LGL: large granular lymphocyte
ERV: endogenous retrovirus
HERV: human endogenous retrovirus
HERV-K: human endogenous retrovirus type K
KGP: 1000 genomes project
HTLV: human T-lymphotropic virus
WGS: whole genome sequencing

## Additional files

**File S1**. Supplementary Methods and Data. Figure S1-S4 are embedded.

**Table S1**. Results of 10 simulated runs to determine how the proportion of infected cells and the length of the retrovirus affects detection by the insertion pipeline.

**Table S2**. The output of the pipeline to investigate HERV-K proviruses and prevalence of polymorphic HERV-K proviruses in 51 LGL leukemia patients.

## Declarations

### Ethics approval and consent to participate

Samples were obtained and informed consents signed for sample collection according to the Declaration of Helsinki using a protocol approved by the Institutional Review Board of the University of Virginia.

### Consent for publication

Not applicable

### Availability of data and material

The complete WGS datasets analyzed during the current study are not publicly available because a full analysis of the genome sequence for other purposes is ongoing. However, the bam files used for analysis of HERV-K provirus occupancy will be made publically available is the manuscript is accepted for publication. The long insert mate pair data from 11 patients are available from the corresponding author on reasonable request.

### Competing interests

The authors declare no competing interests.

### Funding

This research was funded by the National Cancer Institute of the National Institutes of Health under award number R01CA178393, R01CA170334 and P30CA044579 (T.P.L.). The content is solely the responsibility of the authors and does not necessarily represent the official views of the National Institutes of Health. Additional funding was provided to T.P.L. by the Bess Family Charitable Fund, the LGL Leukemia Foundation and a generous anonymous donor. W.L., L.Y. and R.H. were supported in part by R01CA170334. W.L. was also supported in part by the Louis S. and Sara S. Michael Endowed Graduate Fellowship in Engineering and the Fred A. and Susan Breidenbach Graduate Fellowship in Engineering.

### Authors’ contributions

WL developed the HERV-K analysis tool [17], and applied it to the LGL patient dataset. LY implemented the retrovirus search tools used in the insertion call pipeline and the individual read analysis for retroviruses. RH created the insertion call pipeline and conducted simulations. LL implemented the statistical analysis for the HERV-K pipeline tool, TLO supervised the Illumina sequencing and the empirical confirmation of polymorphic HERV-K integration sites; CEH conducted the PCR to confirm polymorphic HERV-K integration sites; DJF oversaw the sample collection and provided critical comments on the manuscript, TPL provided critical discussion on project design and on the final manuscript and secured funding; MP supervised the research, analyzed data and wrote the paper. All authors read and approved the final manuscript.

## Acknowledgments

LGL leukemia patient samples and clinical information were obtained from the LGL Leukemia Registry at the University of Virginia with the assistance of Holly Davis, Bryna Shemo and Andrea Hines. Alexander Wendling, Matthew Schmachtenberg and Shubha Dighe provided excellent technical support while processing patient samples. We thank Aakrosh Ratan for supplying information on the ethnic origin of the 51 LGL patients based on WGS data and for implementing the data mining of the Illumina WGS data. We thank Dr. Stephan Schuster for furnishing paired end sequence data for LGL leukemia patients S9-S11.

