## Supplementary material for "Retrovirus insertion site analysis of LGL leukemia patient genomes": File S1

**Supplementary Methods**

**Pipeline for retrovirus insertions**

Reads are mapped to the hg19 reference genome in two stages.  The first stage uses bwa mem [1] with default settings.  The second stage filters those to create a set of high-confidence mappings based on mapping quality and length (QL-filtering).

QL filtering is accomplished by discarding any mates that fail any of these conditions:

1. Mapping quality of less than 40.  Mapping quality is the bam file’s MAPQ field, as computed by bwa mem.  MAPQ < 40 is supposed to indicate that the probability of being incorrectly mapped is worse than 10^-4^.
2. Less than 40% of the mate is mapped.
3. Both ends of the mate have more than 5 unmapped bases (i.e. we reject mates that only map in the middle).

Insertion events are called as a logical combination of indicator tracks.  Indicator tracks indicate events in corresponding signal tracks.  Signal tracks are derived from the mapping data.  Indicator tracks are two-valued, 1 and 0, while signal tracks are either integer- or real-valued.

A small sample of the reads (about 1 million pairs) is used in a preliminary step to determine the distribution of insert lengths. Pairs in which both mates map within 20K bp with proper orientation provide an estimate of the distribution. The distribution is manually expected to confirm the intended “known” distribution from the physical sequencing experiment. This step is performed separately for each input set.

The distribution is then used to set length thresholds to partition pairs into three classes — short, normal, and long.  The “normal” class consists of lengths roughly between the 5% and 95% percentiles of the distribution.

Step 1. Reference genome blacklisting indicator tracks

Two blacklisting tracks are created to indicate regions where mapping depth can be expected to be inconsistent with sequence content.  One track indicates unassembled regions in the reference, N-runs.  The other track indicates areas of known repeats. These two tracks are used in the pipeline to counteract negative effects in signal computation in the vicinity of such elements.

The reference genome N-runs indicator track is derived from the reference genome (hg19) sequence.  The track consists of all runs of 50 or more consecutive Ns. The result is a two-valued track, values 1 and 0, with 1 indicating positions that are part of a run of Ns.

The Repeat Masker indicator track is derived from the hg19 Repeat Masker file, rmsk.txt.gz, at the USCS genome browser.  All intervals from the Repeat Masker file are copied, merging any overlaps.  All other annotations are discarded.

Step 2. Average mate pair insert length signal track, and indicator track [Track 1]

The average MP insert length *signal* track is derived from the bam files of the QL-filtered long insert mate pair WGS data of LGL.  Reads in which both mates map in the proper orientation (tail-to-tail) and with a plausible insert distance (between 300 and 30K bp) are identified.  At each genomic position, the average insert length of the reads covering that position is computed.  A read covers a position if the position falls with the mapped interval of either mate in the read, or in the interval between them.  The result is a real-valued track showing the average *inferred* insert length over the genome.  To facilitate viewing, the median insert length is subtracted from this track.

Where no structural changes have occurred we expect this track to be consistent with the average length of the insert molecules (track value near zero).  Variation from this average indicates insertions or deletions in the subject relative to the reference genome.  For example, an insertion in the subject should be apparent as a decrease in the insert length track.

The average mate pair (MP) insert length *indicator* track is derived from the average MP insert length signal track.  Conceptually we are looking for troughs in the signal.  This is accomplished by identifying intervals of length at least 500 bp in which the signal averages no more than the 5th percentile.  Specifically, the distribution of the average of the signal in 500 bp windows is estimated, and the 3rd, 5th, and 60th percentiles are determined.  The signal is clipped at the 3rd percentile (values less than the threshold are changed to the threshold) after which intervals meeting the 5th percentile and length criteria are identified.  The resulting track is 1 within such intervals, and 0 outside it.  The signal clipping step is motivated by the observation that very deep troughs widen the reported intervals, with the potential to blur two events together.  Additionally, since long unassembled regions and repeats in the reference genome generate false low signal and confuse this computation, the signal is first masked by setting all blacklisted intervals to the value of the 60th percentile.  Additionally, all blacklist intervals are masked to zero in the result track.

Step 3. Mate pair short and normal insert coverage depth signal tracks [Track 2 & 3, respectively], and indicator tracks

Two MP insert coverage depth *signal* tracks are derived from the bam files of the QL-filtered long insert mate pair WGS data of LGL.  Reads in which both mates map in the proper orientation (tail-to-tail) and with insert distance within the corresponding range (300bp to 6Kb for short or 6Kb to 10Kb for normal) are identified.  At each genomic position, the number of reads covering that position is counted.  A read covers a position if the position falls within the mapped interval of either mate in the read, or in the interval between them.  The result is two integer-valued tracks showing the coverage depth over the genome for each class of insert lengths.

Where no structural change has occurred we expect nearly all reads to have a normal insert length; the short track should have no depth while the normal track should have depth consistent with the average sequencing depth.  Variation from this condition indicates insertions or deletions in the subject relative to the reference genome.  For example, an insertion in the subject should be apparent as an increase in the depth of short inserts and a decrease in the depth of normal inserts.

The normal insert coverage depth *indicator* track is derived from the normal insert coverage depth *signal* track.  Conceptually we are looking for troughs in the signal.  This is accomplished by identifying intervals of length at least 500 bp in which the signal averages no more than the 5th percentile.  Specifically, the distribution of the average of the signal in 500 bp windows is estimated, and the 5th and 60th percentiles are determined.  Intervals meeting the 5th percentile and length criteria are identified. The resulting track is 1 within such intervals, and 0 outside it. The blacklisted intervals are incorporated the same as for the average MP insert length indicator track.

The short insert coverage depth *indicator* track is derived from the short insert coverage depth *signal* track.  Conceptually we are looking for peaks in the signal.  This is accomplished by identifying intervals of length at least 1,000 bp in which the signal averages no less than the 90th percentile.  Specifically, the distribution of the average of the signal in 1,000 bp windows is estimated, and the 90th (and 92nd) percentiles are determined.  The signal is clipped at the 92nd percentile (values greater than the threshold are changed to the threshold) after which intervals meeting the 90th percentile and length criteria are identified.  The resulting track is 1 within such intervals, and 0 outside it.  The signal clipping step is motivated by the observation that very tall peaks widen the reported intervals, with the potential to blur two events together.

Step 4. Mate pair discordant mates coverage depth signal track, and indicator track. [Track 4]

To clarify the terms used in this section, a pair of discordant mates means that the read pair is either broken (only one mate of the pair mapped), bichromosomal (the mates map to different chromosomes) or distant (the mates map to the same chromosome but are more than 30kbp away from each other).  An anchoring mate refers to the mate of the read mate pair that mapped with the quality described above to a given genomic location on hg19 and a discordant mate refers to the mate of the read pair that is either unmapped or mapped to a distant or bichromosomal location other than the given genomic location.

The mate pair discordant mates coverage depth signal track is derived from the bam files of the QL-filtered long insert mate pair WGS data of LGL.   Anchoring discordant mate pairs are extracted from the bam file and filtered with mapping quality >= 40 and at least 40% of the sequence mapped to stay consistent with the overall mapped read filter scheme; the discordant mates are additionally trimmed for average MAPQ >= 20 at the ends and at least 60bp after trimming.  A discordant mate pair is only kept if both its anchoring and discordant mate pass the quality filter.  At each genomic position, the number of anchoring mates covering that position is counted.  An anchoring mate covers a position if the position falls with the anchoring mate’s mapped interval.  The result is an integer-valued track showing the coverage depth of discordant mates over the genome.

The discordant mates coverage depth *indicator* track is derived from the discordant mates coverage depth *signal* track.  Conceptually we are looking for clumps of higher occupancy in the signal.  This is accomplished by identifying intervals of length at least 8,000 bp in which the signal’s occupancy is no less than the 95th percentile.  Specifically, distribution of the occupancy of the signal in 8,000 bp windows is estimated, and the 95th percentile is determined.  Intervals meeting the 95th percentile and length criteria are identified.  The resulting track is 1 within such intervals, and 0 outside it.  Occupancy is a count of the number of non-zero positions in the window-- positions covered by at least one discordant broken mate.  Since this is equivalent to the average of the binarized signal over the window, the implementation binarizes the signal before performing the computation.

Step 5. Paired end clipped breakpoints signal track, and indicator track [Track 5]

The PE clipped breakpoints *signal* track is derived from the bam files of the QL-filtered paired end WGS data of LGL.  Mates that the aligner has mapped only in part are identified (“clipped mates”).  Such mates in which the unmapped portion is shorter than 10% of the mate’s length are discarded (presumed to be the result of sequence quality issues).  Each remaining mate suggests a single location in the genome where a rearrangement breakpoint has occurred.  At each genomic position, the number of mates suggesting breakpoints at that position is counted.  Mates suggesting breakpoints beyond the end of the reference sequence are discarded (this is more prevalent with certain reference contigs).  This intermediate result shows the number of times a particular breakpoint has been suggested.  Since aligners typically do not have single base accuracy for breakpoints, the intermediate result is passed through a 10-bp sliding sum.  The result is an integer-valued track showing the number of times any breakpoint has been suggested within ±5 bp, over the genome.

Where no rearrangement events have occurred we expect there to be no suggested breakpoints.

The clipped breakpoints *indicator* track is derived from the clipped breakpoints signal track.  Conceptually we are looking for positions of high value in the signal. This is accomplished by identifying locations where the signal’s value is no less than the 96.2th percentile.  Specifically, distribution of the signal’s value is estimated, and the 96.2th percentile is determined.  The signal is binarized at that percentile, so that the resulting track is 1 at positions at or above the threshold, and 0 otherwise.

Step 6, insertion calls track

The insertion calls signal track is derived from the five indicator tracks created in the preceding steps as described above. A summary of the tracks is as follows (Figure S1). The first four tracks all relate to mate pair mapping and track 5 is derived from short insert paired end sequence data.

Track	Description

1	average mate pair insert length indicator track

2	normal insert coverage depth indicator track

3	short insert coverage depth indicator track

4	discordant mates coverage depth indicator track

5	clipped breakpoints indicator track

Conceptually, the insertion calls indicator track can be represented as a simple logical combination of the indicator tracks.

(Track 1 or Track 2) and (Track 3 or Track 4) and Track 5

In that description “or” represents the usual binary operation, with the result having a 1 wherever either input was 1, and zero only where both inputs were zero.

However, the “and” allows for signals that occur proximally without actually overlapping. Indicators are considered to overlap if they are within ±2000 bp.  Effectively we first identify candidate intervals with (Track 1 or Track 2) then use the other indicator tracks as filters.  If a candidate does not have any proximal interval in either of (Track 3 or Track 4) it is discarded.  Similarly, if it does not have any proximal interval in Track 5, it is discarded.  Candidates surviving these two filter steps comprise the insertion calls track.  However, because Track 1 or Track 2 tend to be highly fragmented by blacklist masking, a final merging step is applied to remove all between-interval gaps of less than 2000 bp.

**Insertion Simulation**

A simple simulated data set was generated in order to guide the setting of parameters in the insertion pipeline. The simulation represented non-interfering insertion events of lengths ranging from 1Kbp to 12Kbp with infection rates from 50% to 100%. As described below, the generation of simulated reads was guided by the lengths and sequencing depths of a real subject (S10). The ability of the pipeline to detect, or not detect, these events guided the manual setting of pipeline parameters with a goal of being able to detect insertions with length ranging from 6-10kb in the presence of infection rates of at least 50%.

The simulation modeled the reference genome as a 11Mbp randomly generated sequence of ACGT, with no nucleotide bias. An uninfected “subject” genome was generated with 1% divergence from the reference (substitutions only, i.e. any location had a 1% chance of being different than the reference). An infected subject genome was generated from the subject by the insertion of random sequences. These “insertion events” occurred at the midpoints of non-overlapping 100Kbp windows. Insertions had specific lengths -- 1Kbp, 3K, 4K, 5K, 6K, 7K, 8K, 9K, 10K and 12K -- and the distance of 100K between events was long enough to ensure that every event would behave independently in the pipeline. To facilitate study of various infection rates, this process was repeated eleven times to create eleven different 1Mbp subunits, each with ten insertion events.

Sequenced reads were modeled (“generated”) by pulling random intervals from the infected and uninfected genomes, with an error rate of 1% (substitutions only). Two sets of reads were generated to match the data available for the real subject.  For paired end, reads of length 101x101 with insert distribution centered at 300 bp were generated to 30X coverage depth.  For mate pairs, reads of length 150x150 with insert distribution centered at 8K bp were generated to 5X coverage of the sequences, which corresponds to ~133X coverage including the unsequenced insert.

Read generation was performed independently for each of the eleven subunits. The subunits corresponding to infection rates ranged from 50% to 100% in 5% increments. In each subunit, the ratio of reads pulled from the infected subject versus uninfected was controlled by the infection rate for that subunit.

The reads for all subunits were mixed together and mapped to the reference.  Since the reference genome was randomly generated the probability of mismapping is effectively zero. Reads generated for a particular subunit will only map to the neighborhood corresponding to that subunit (or not map). Though all reads are processed together in the pipeline, in effect they are a different experiment for each insertion length and infection rate.

This simulation is intentionally naive, focusing on insertion length and infection rate while eliminating real world factors such as repeats, nucleotide bias, deletions, short indels, divergence rates that vary between content (e.g. coding vs non-coding), unassembled intervals in the reference, locational sequenceability bias, and others. The number of detections in 10 simulated runs for inserts at different lengths and levels of infection is shown in Table S1. An example of the output for an infection rate of 75% is shown in Figure S1.

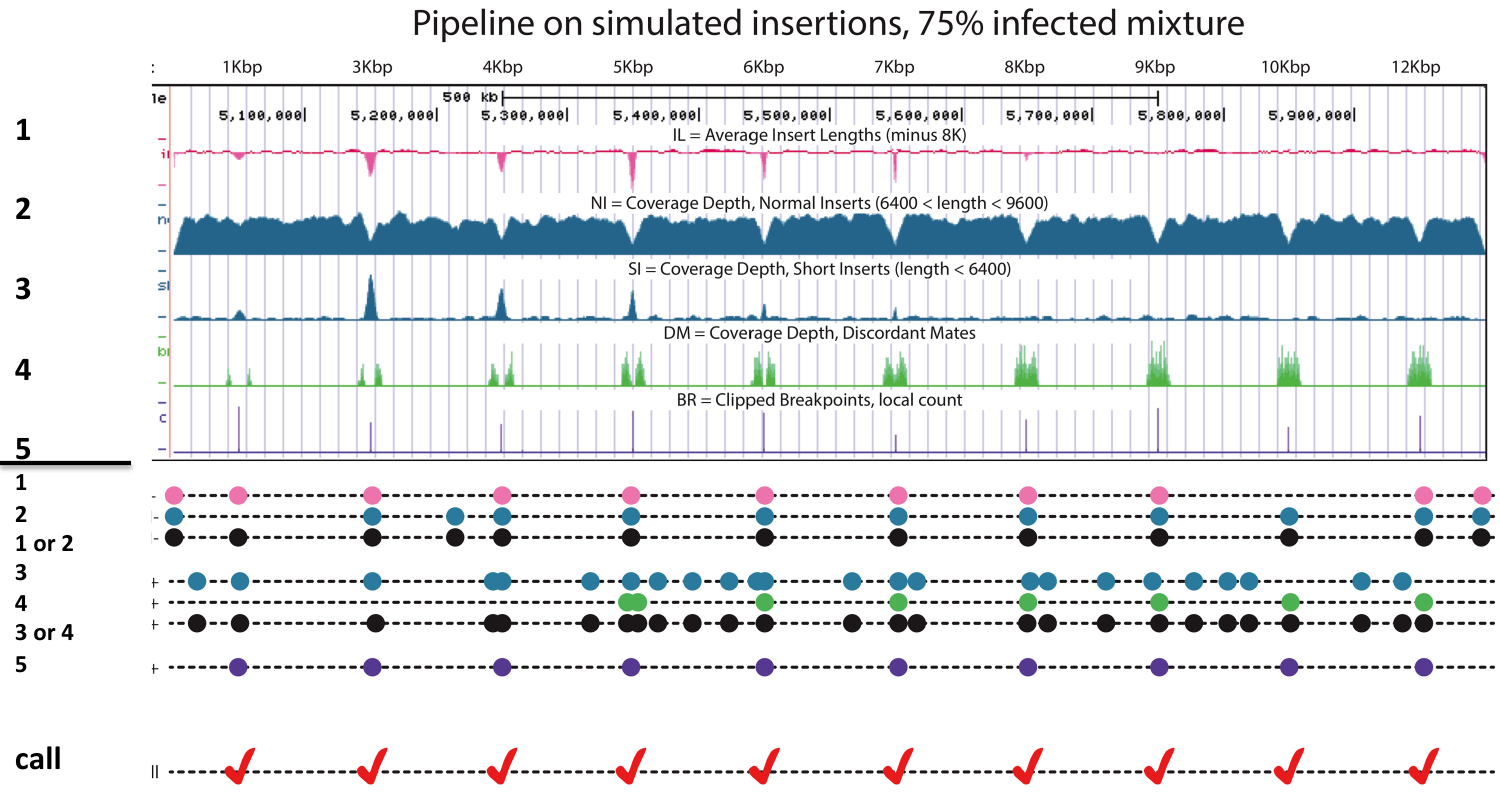

**Figure S1. An example of the tracks used to detect an insertion of varying length when the proportion of infected cells is 75%. In the upper panel, tracks are numbered as in text.** 1: average insert length of mapped mate pair reads [subtracted from 8000 so the number is negative for short inserts], 2: depth of mate pair reads with expected insert length, 3: depth of mate pair reads with shorter than expected insert length, 4: depth of anchoring discordant mate pair, 5: depth of clipped (partially mapped) short insert paired-end reads. Note how the signal for tracks 1 and 3 is lost at longer insert length. However, the signal requires either track 1 or 2, and either track 3 or 4 to provide a signal to call an insertion, as shown in the lower panel and described in the text.

The parameters determined by the simulation were applied to the long insert mate pair data of 11 LGL leukemia patients. An example of the tracks generated at an insertion call is shown in Figure S2.

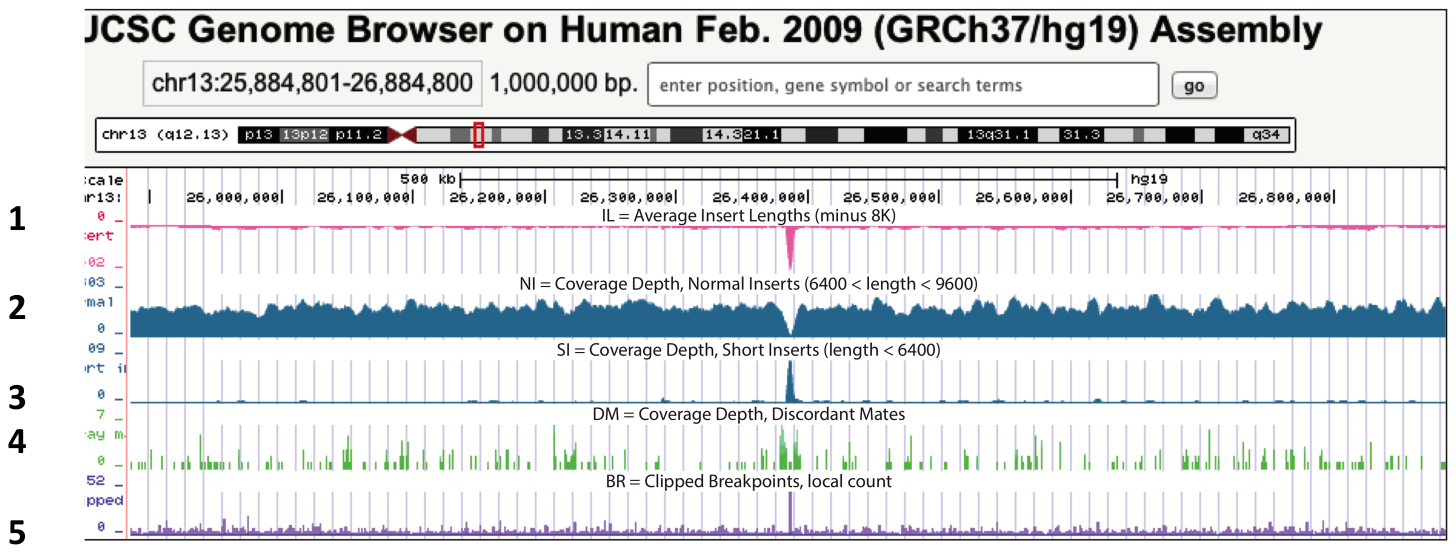

**Figure S2. Example of a signal track for an insertion call from sequence data (patient S09).** The numbers on the left side indicate track codes as described in supplementary methods and Figure S1. In this case the insertion site is clearly defined by all tracks.

**Detection of retrovirus in the insertion call pipeline**

Each candidate genomic insertion call for the 11 LGL leukemia patients with long insert mate pair WGS was next evaluated for candidate clonal retrovirus insertions (Figure S3A).   Sequences were retrieved from the discordant mate pairs around each insertion call: mate pair reads whose mate anchor at +-30 kb of the insertion were collected using a custom perl script that parses the bam file.  The collected reads included the discordant reads that mapped to a distant location of the same chromosome and reads that mapped to another chromosome.  Extracted reads at each insertion site were assembled with cap3 [2].  Cap3-assembled contigs were searched against the NCBI nt database (downloaded on July 2nd, 2016, this version of the nt database is used for all analyses described) using BLAST+ [3, 4].  BLAST hits were sorted by the e-value of the alignments and tagged for having human or retrovirus taxonomy.  Human hits were labeled for having taxonomy ID 9606, and retrovirus taxonomies were identified with a custom perl script that starts with the taxonomy ID of the BLAST hit and iteratively searches the NCBI taxonomy ID table from the tip of the tree until reaching the node named “Retroviridae”.  A candidate infectious retrovirus insertion features at least two contigs whose lowest e-value BLAST hits have Retroviridae taxonomy.  Similarly, a candidate human endogenous retrovirus (HERV) insertion features at least two contigs whose lowest e-value BLAST hits have taxonomy ID 9606 and “endogenous retrovirus” or “ERV” in its entry name.  For HERV candidates, fosmid or BAC (bacterial artificial chromosome) hits were mapped back to the genomic location around the insertion call, and a reference HERV insertion sequence was mapped to the fosmid or BAC sequence, both done with blat [5].  A HERV insertion candidate is only called when the fosmid or BAC can be mapped back to the genome at full length and high identity with an alignment gap where the HERV reference maps to the fosmid or BAC.

**Alternative analysis for new HERV-K insertions**

Our insertion call pipeline requires that over 55% of cells in a sample contain the integrated retrovirus, a condition that would be met under the assumption that the retrovirus integrated prior to the clonal expansion of the LGL cells. We also utilized the long insert mate pair whole genome sequencing data to identify HERV-K proviral insertions in the 11 LGL patients that could appear as a somatic integration subsequent to the expansion of the LGL cells or in non-cancer cells in the sample.  This approach is neither dependent on the known HERV-Ks in the reference genome, nor on the *de novo* insertion calls presented in this paper. Long insert mate pair reads were mapped to hg19 (downloaded from http://hgdownload.cse.ucsc.edu/goldenPath/hg19/bigZips/hg19.2bit) and the coding part of a full-length HERV-K reference (JN675087 from GenBank) separately using bwa mem [1] default parameters.  Mapped sequences were sorted and compressed using samtools [6] default parameters.  Mate pairs with one read mapped to a non-HERV-K location of hg19 and the other read mapped to JN675087 were extracted using a custom perl script and trimmed to remove adapters and low quality ends using trimmomatic [7] under the following parameters: PE ILLUMINACLIP:TruSeq3-PE.fa:2:30:10 LEADING:3 TRAILING:3 SLIDINGWINDOW:4:15.  The reads of the pair that map to hg19 were clustered using bedtools cluster [8] requiring reads in the cluster to be mapped on the same strand of the genome (parameter “-s”) and allowing maximum 1,497 bp gap (parameter “-d 1497”) between reads.  Given the nature of Illumina Nextera long insert mate pair sequencing libraries, a HERV-K in hg19 corresponds to a pair of such anchoring read clusters: one on the reverse complement strand upstream of the HERV-K, and another one on the sense strand downstream of the HERV-K.  The “-d 1497” parameter was chosen to maximize the number of reads in all the anchoring read cluster pairs.  Anchoring read cluster pairs are able to identify HERV-K either assembled in the hg19 reference or not: cluster pairs that are distant from a known HERV-K location in hg19 are used to identify novel HERV-K insertions; mates of the reads in the anchoring read cluster pairs within 30 kb of a known HERV-K location were mapped back to the HERV-K reference at that location as a validation of the approach.

**Rare retrovirus sequence detection**

We thoroughly analyzed each unmapped and unassembled read that passed the read quality filter for a rare retrovirus insertion (Figure S3B). These reads include unmapped mate pair and paired end WGS reads of 11 patients [S1-S11].  We first assembled all collected reads using minia (https://github.com/GATB/minia, for WGS) default parameters. Reads that cannot be assembled have their low quality ends and adapters trimmed off using trimmomatic under the following parameters: PE ILLUMINACLIP:Nextera-MP.fa:2:30:10 LEADING:3 TRAILING:3 SLIDINGWINDOW:4:15.  Inspired by the ideas of Naccache *et al.* [9], trimmed unassembled WGS sequences and singlet contigs were aligned to the NCBI nt database (downloaded on July 2nd, 2016, this version of the nt database is used for all analyses involving it) with SNAP [10] to quickly assign a taxonomy ID using the parameters of “-locationSize 5”, “-sc” 1 and “-pc”.  Non-vertebrate taxonomy ID hits were identified by an iterative search of the NCBI taxonomy ID table starting from the tip of the tree until reaching the node named “nonvertebrata” using a custom perl script.  Two batches of reads went through a slower but more sensitive query using BLAST.  These include the minia-assembled contigs of the unmapped WGS reads (Figure S3B, second row, left), the unmapped, unassembled and non-vertebrate-taxonomied reads (Figure S3B, third row, right).  These sequences were aligned to the virus database in the NCBI nt database using blastn of the NCBI BLAST+ software to identify any potential retroviral sequences.  The virus database was extracted with a custom perl script that iteratively search the NCBI taxonomy ID table from the tip of the tree until reaching the node named “Vira”.  Endogenous retroviruses were also included in the virus database by searching with retrovirus-related keywords in the name of the nt database entry and manual curation.  Any entry with a “Vira” taxonomy ID or that contains an endogenous retrovirus keyword in the name is extracted from the NCBI nt database to be included in the virus database.  BLAST hits against the virus database were sorted by their e-value, and only the sequences with a best match to an infectious or endogenous retrovirus (represented by the lowest e-value of any match) were considered as a candidate rare retrovirus. Finally, the reads or contigs of candidate retroviruses were re-submitted to BLAST against the whole nt database, and only the ones whose unique top hit were still retrovirus were considered candidate rare retrovirus insertions.

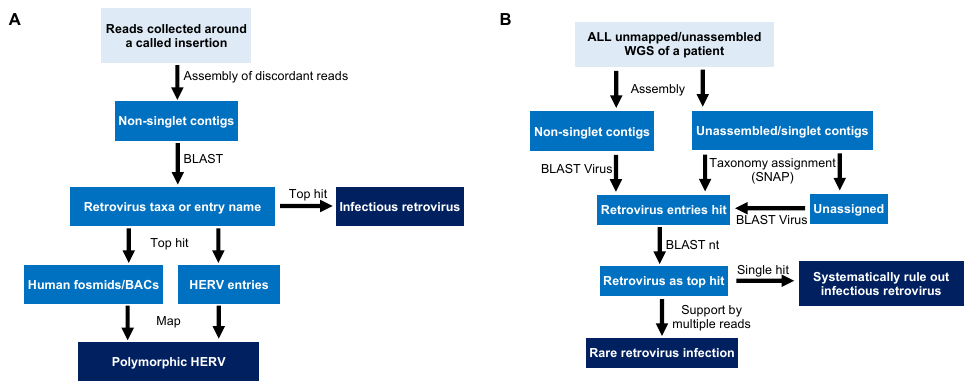

**Figure S3. Diagram of retrovirus identification pipeline.** Data and results are shown in boxes and steps and conditions taken are shown by arrows and text next to them. **A.** Pipeline used for the clonal retrovirus insertion sequence analysis as described in Supplementary Methods “Detection of retrovirus in the insertion call pipeline”. Detection is dependent on the assumption that the retrovirus integrated prior to the clonal expansion of LGL cells and is present in more than 55% of cells in the sample. **B.** Pipeline for rare retrovirus detection. This approach was taken to detect a retrovirus integration that was present in a small proportion [less than 55%] of cells in the sample.

**Supplementary Data**

Figure S4 demonstrates the relationship between intermediate *n/T* vales and HERV-K alleles.

**A**

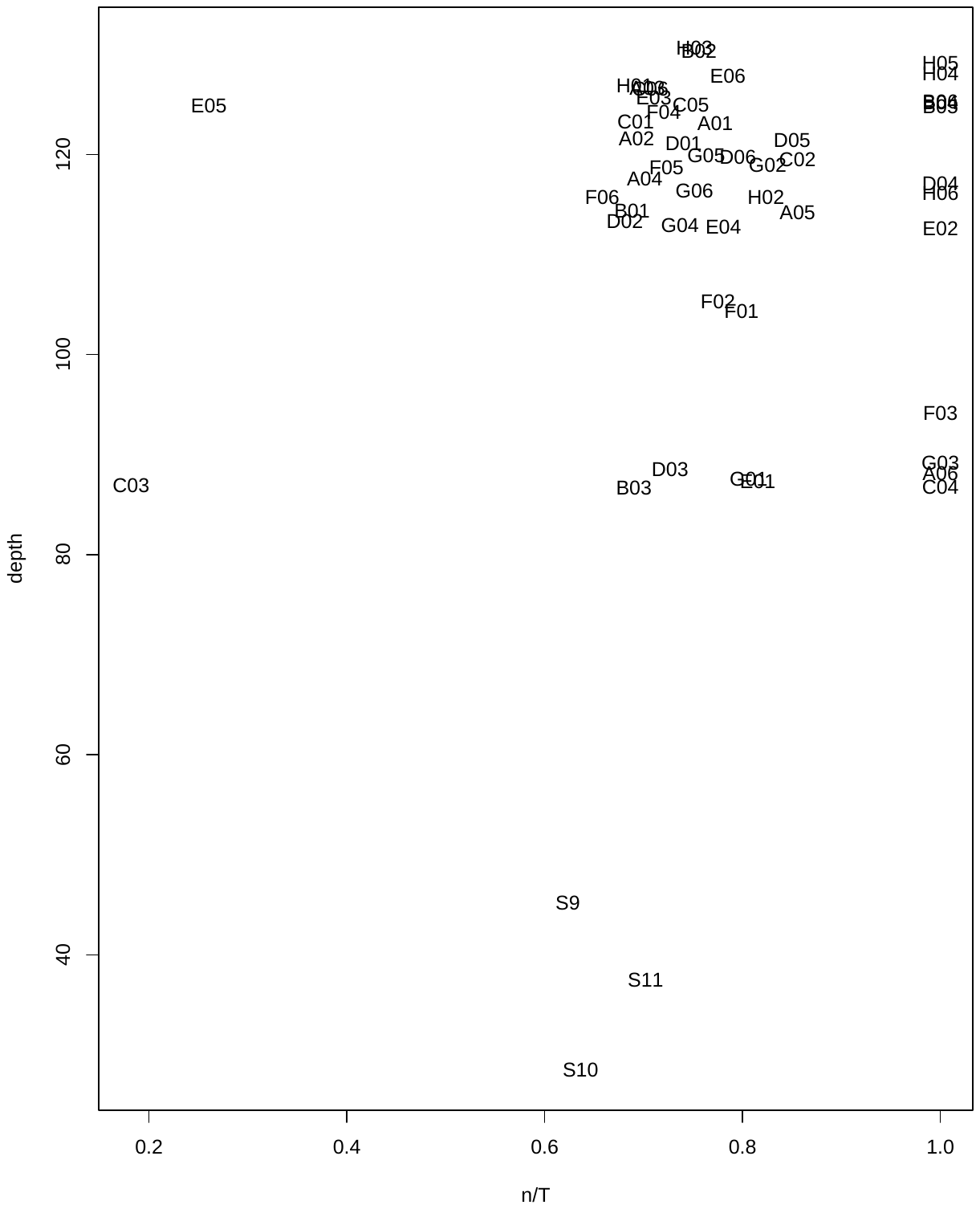

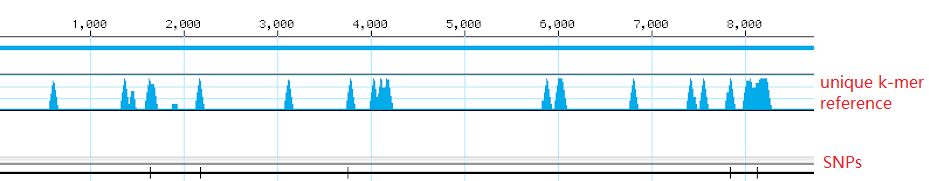

**B**

**Figure S4. Exploration of n/T value and sequence polymorphism for chr3:112743479-112752282 (hg19) in LGL leukemia data.** **A.** The n/T vs. depth plot for 51 LGL leukemia patients. The x-axis reflects the n/T ratio, which indicates the proportion of unique sites for a specific HERV-K (T) identified in the patient sequence data (n). Y-axis is sequence depth. In the plot the majority of patients do not have the reference allele [n/T= 1] for this HERV-K but there is considerable variation in the n/T values suggesting that several alleles could be present. Note two patients (C03, E05) have a solo LTR [n/T ~0.2, confirmed by mapping k-mers on reference HERV-K for this locus]. **B.** Depiction of SNP locations in the chr3:112743479-112752282 HERV-K sequence from LGL leukemia patients. The bar at the top represents the coordinates of the HERV-K and the blue peaks show the location of k-mer set T [all k-mers unique to this HERV-K] for this virus. SNPs are depicted at five positions indicated by black lines on the bottom line. A SNP must overlap with a unique k-mer from set T to impact the n/T ratio.
